# Integration and modularity in Procrustes shape data: is there a risk of spurious results?

**DOI:** 10.1101/371187

**Authors:** Andrea Cardini

## Abstract

Studies of morphological integration and modularity are a hot topic in evolutionary developmental biology. Geometric morphometrics using Procrustes methods offers powerful tools to quantitatively investigate morphological variation and, within this methodological framework, a number of different methods has been put forward to test if different regions within an anatomical structure behave like modules or, *vice versa*, are highly integrated and covary strongly. Although some exploratory techniques do not require *a priori* modules, commonly modules are specified in advance based on prior knowledge. Once this is done, most of the methods can be applied either by subdividing modules and performing separate Procrustes alignments or by splitting shape coordinates of anatomical landmarks into modules after a common superimposition. This second approach is particularly interesting because, contrary to completely separate blocks analyses, it preserves information on relative size and position of the putative modules. However, it also violates one of the fundamental assumptions on which Procrustes methods are based, which is that one should not analyse or interpret subsets of landmarks from a common superimposition, because the choice of that superimposition is purely based on statistical convenience (although with sound theoretical foundations) and not on a biological model of variance and covariance. In this study, I offer a first investigation of the effects of testing integration and modularity within a configuration of commonly superimposed landmarks using some of the most widely employed statistical methods available to this aim. When applied to simulated shapes with random non-modular isotropic variation, standard methods frequently recovered significant but arbitrary patterns of integration and modularity. Re-superimposing landmarks within each module, before testing integration or modularity, generally removes this artifact. The study, although preliminary and exploratory in nature, raises an important issue and indicates an avenue for future research. It also suggests that great caution should be exercised in the application and interpretation of findings from analyses of modularity and integration using Procrustes shape data, and that issues might be even more serious using some of the most common methods for handling the increasing popular semilandmark data used to analyse 2D outlines and 3D surfaces.

## Modularity, Integration, and the ‘Procrustes Paradigm’ in Geometric Morphometrics

The analysis of modularity and integration using geometric morphometrics methods based on Procrustes shape coordinates has become increasing popular in evolutionary developmental biology (Mitteroecker and Bookstein 2007;Klingenberg 2009, 2013a, 2014). Simplifying, the main question asked in this type of studies is whether morphology varies during ontogeny and evolution as a single highly integrated structure or as an ensemble of more or less loosely correlated subunits (called modules); in this second case, the covariation within parts of the module is expected to be higher than among parts of different modules. For instance, one might assess the strength of the covariation between the face, vault and cranial base during cranial ontogeny in humans (Bookstein et al. 2003): if low, that means that these anatomical regions might change relatively independently (of course, within reasonable limits) and thus behave like ‘modules’; if high, it implies that cranial shape varies as whole in a highly coordinated fashion, so that changes in one region are accompanied by modifications in all the others, and, in theory, one can predict changes in the face or vault by knowing what happens in the cranial base or *vice versa* (Klingenberg 2013a).

Landmark-based geometric morphometrics employs Cartesian coordinates of corresponding anatomical points, called landmarks, to quantify size and shape (or form, when size and shape are put back together -Mitteroecker et al. 2013) of different individuals, ontogenetic stages, species etc. (Adams et al. 2004, 2013;Cardini 2013). Shape is obtained from the original raw coordinates by standardizing size and minimizing translational and positional differences in a sample. This is generally achieved using a generalized Procrustes analysis (GPA -Rohlf and Slice 1990), which is a least square procedure that, after dividing the raw coordinates by their centroid size, centers all observations at their centroids and iteratively rotates them to minimize the sum of squared distances to the sample mean.

The Procrustes superimposition, however, is not the only method to obtain shape coordinates from Cartesian coordinates of landmarks. Also, a GPA is clearly not based on any biological model of developmental and evolutionary variation, as it is simply a least square approach like, for instance, that used in an ordinary linear regression to find an approximation for the relationship between a predictor and a dependent variable. To make a crude analogy using molecular biology as an example, one can think of the Procrustes superimposition as akin to devising an alignment of DNA sequences that simply maximizes the correspondence of all bases by weighting them all equally. However, this is not what molecular biologists normally do, because we have a good understanding of molecular evolution and know that changes in certain positions are more likely to occur (e.g., the third base of a codon), deletion or insertion may be more frequent in less conserved genes, some DNA regions evolve more slowly than others etc. Thus, sophisticated alignments can be tried, which take our knowledge of molecular evolution into account (Felsenstein 2004, Chapter 29). For morphology, in contrast, we do not have the same type of deep understanding as for DNA changes, and we have therefore adopted the ‘Procrustes paradigm’ (Adams et al. 2013) to minimize positional differences, because of its desirable statistical properties and generally better performance compared to alternative methods (Rohlf 2000a b;, 2003;Adams et al. 2004). It is thus important to always bear in mind that Procrustes, by far the most used method in landmark-based geometric morphometrics, is a very convenient and mathematically well defined (Rohlf 2000ab;, 2003) but biologically arbitrary choice to derive shape coordinates. This is true also for other superimposition methods, such as Bookstein’s baseline or resistant fit (Rohlf and Slice 1990;Bookstein 1991). Indeed, most of the issues raised by this study apply to those methods as well. In fact, these issues do not detract from the demonstrated merits of Procrustes, that led to the current paradigm in geometric morphometrics (Adams et al. 2013). However, raising them, although in the context of a set of very specific types of analyses, might help to recall some of its limitations, that have largely been acknowledged since the early days of geometric morphometrics by the leading figures of this field, as nicely reviwed byO’Higgins (2000).

## Approaches to Modularity and Integration using Procrustes Shape Data

In studies of modularity and integration one has a battery of potential tests that can be employed on Procrustes shape data (Mitteroecker and Bookstein 2007; Marquez 2008;Klingenberg 2009, 2013a;Goswami and Polly 2010;Fruciano et al. 2013;Adams 2016;Adams and Collyer 2016;Goswami and Finarelli, 2016). However, as nicely reviewed by Baab (2013), and in more detail by Klingenberg (2009,2013a), when *a priori* modules are part of a single structure, such as, for instance, the cranium, before any test is done, one has first to take a decision between two different approaches. The first is sometimes (Klingenberg 2011) called the ‘within a configuration’ approach. In this case, a morphometrician does a single Procrustes superimposition including all landmarks, obtains a unique set of shape coordinates and then splits these shape coordinates into modules made of different subsets of landmarks. Alternatively, using the ‘separate blocks’ approach (Klingenberg 2011), one can start with the raw data (no superimposition), split them into modules and then perform separate Procrustes superimpositions for each module.

The difference between the two approaches is substantial, because, in the first case, the relative size and positional differences of all modules are preserved, with all data being in the same common shape space (Bookstein et al. 2003; Klinbengerg 2009;Baab 2013). In the second approach, in contrast, each module is in its own separate and unique shape space, while the information on the relative size and position of the modules is discarded. The initial decision of which approach to follow has, therefore, potentially profound influences on results, and one or the other strategy might be more appropriate depending on the specific structure and scientific question (Baab 2013;Klingenberg 2009, 2013a). Nevertheless, it has recently been argued in MORMPHET, the email discussion list of morphometricians, that the within a configuration method might lead to more interpretable results “as with a single GPA one is able to characterize covariation patterns among sets of variables whose spatial relationships have been retained throughout the analysis” (Dean C. Adams, on January 24^th^ 2018). Indeed, as mentioned, using separate blocks, the knowledge about relative sizes and orientation of modules is lost, or only partially preserved if they contain at least a few common landmarks (Klingenberg 2009), thus potentially removing part of their true covariance.

## Why Procrustes Shape Data are ‘Special’

In general, Procrustes shape variables are analysed using multivariate methods, many of which have been borrowed from traditional morphometrics (Neff and Marcus 1980;Marcus 1990). However, Procrustes shape coordinates are ‘special’ because, as anticipated in the previous section, the choice of the method to superimpose specimens and obtain shape is biologically arbitrary (O’Higgins 2000). In practice, this means, for instance, that the coefficients of a principal component analysis (PCA), or those of a multivariate linear regression of shape onto a predictor, cannot be interpreted as in traditional morphometrics, as they depend on the choice of the superimposition. For the same reason, Procrustes shape variables, such as shape coordinates or partial warps (Bookstein 1991), cannot be analysed one at a time (Rohlf 1998). Similarly, patterns of variation of single landmarks from a larger configuration should not be analysed or interpreted after a common superimposition.

In contrast, as long as the full set of multivariate shape variables (all Procrustes shape coordinates or partial warps) are examined, and results, including the visualization, are interpreted by integrating findings over the whole set of landmarks in a configuration, the outcome is correct and generally robust to the choice of the superimposition. This is exemplified in Figure 1 (reproduced from Viscosi and Cardini 2011) using simple triangles. With triangles, one can visualize the overall shape similarity relationships using a scatterplot of the first two PCs, that together summarize 100% of variance. Regardless of whether shape coordinates are obtained using Procrustes or a different method, such as Bookstein’s baseline (1991), which for 2D data rescales, translates and rotates all specimens so that they overlap in two points (the ‘baseline’), the pattern suggested by the scatterplots, as well as the visualization using the thin plate spline (TPS -Bookstein 1991;Klingenberg 2013b) transformation grids, are virtually identical in both cases. In contrast, PC loadings and displacement vectors (which represent shape differences as ‘displacements’ of landmarks in a target shape relative to a reference, such as the sample mean) are radically different using one or the other superimposition method.

**Fig. 1.**
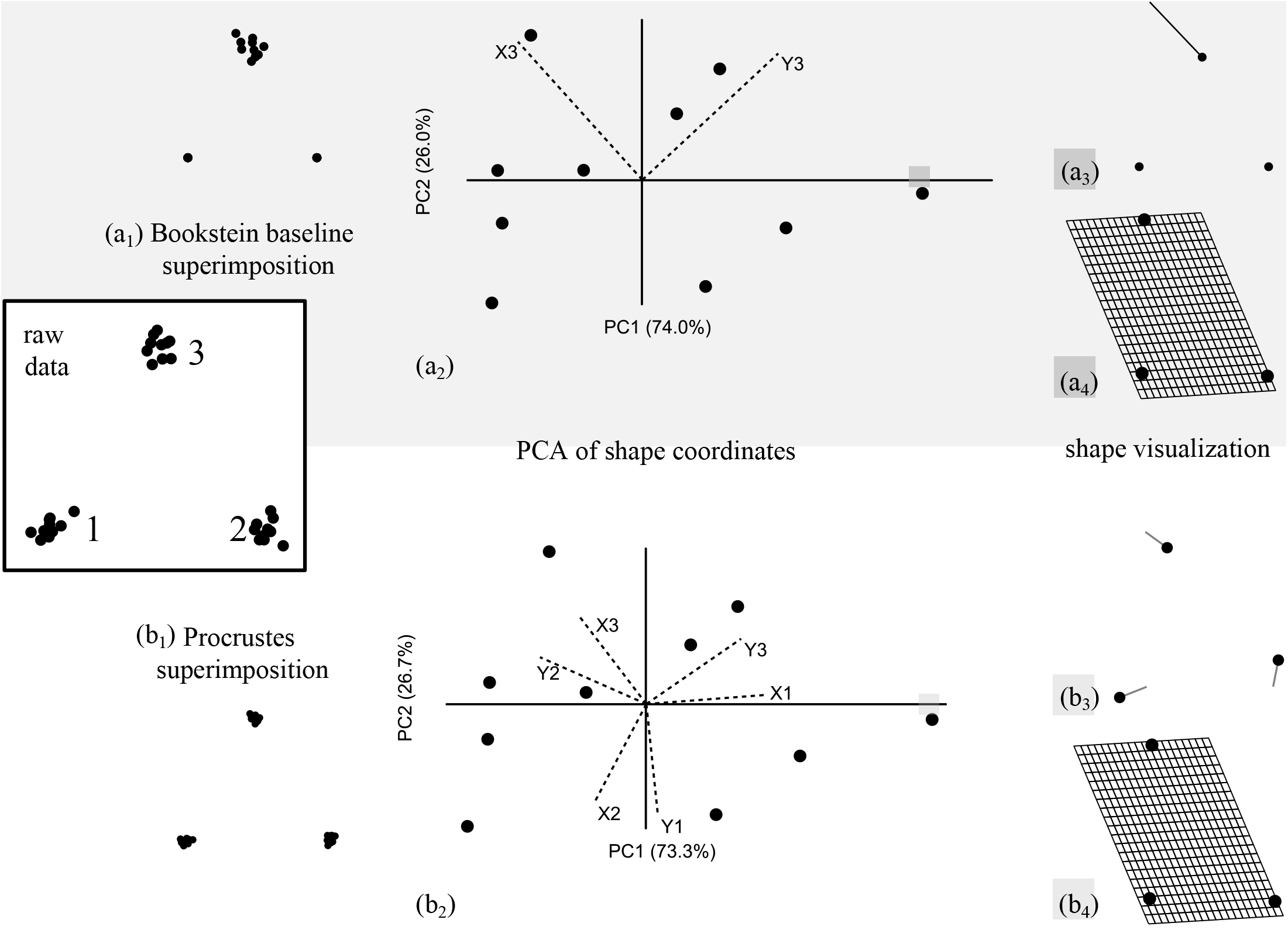
Example of the effect of different superimpositions on the interpretation of results (reprinted from Figure 9 of Viscosi and Cardini 2011, under an open access license: http://journals.plos.org/plosone/s/licenses-and-copyright): PCAs of a sample of 10 random triangles (raw data) superimposed either using Bookstein baseline method (a1, with landmarks 1-2 as a baseline) or Procrustes (b1). Biplots (a2, b2) show the scatterplot of the specimens (filled circles) and the loadings (dotted lines) employed to weight the matrix of shape coordinates (X1, Y1, etc.). As an example, shape variation at the positive extreme of PC1 is visualized (with a four times magnification) using either displacement vectors (a3, b3) or TPS grids (a4, b4).

## Can Procrustes Introduce Covariance that may lead to Spurious Results in analyses of Integration and Modularity within a Structure?

The aim of this study is to explore the consequences of the Procrustes superimposition in analyses of integration and modularity, with a main focus on the popular within a configuration approach. This approach might be particularly problematic, because it makes use of subsets of points after a common superimposition. To frame this issue in a less abstract manner, one can make a comparison with the potentially misleading interpretations of displacement vectors in the visualization of shape differences. In this sense, breaking a set of Procrustes shape coordinates into mutually exclusive subsets of landmarks might be seen as analogous to picking up one displacement vector (or a few) from a bigger configuration and interpreting its(/their) direction as meaningful on its(/their) own, instead of more accurately relating it to the directions of all the other vectors in the entire configuration. For instance, the displacement vector of landmark three, with data superimposed using Bookstein’s baseline, as shown in the upper portion of Figure 1, indicates a sharp upward and backward movement of that specific point. In terms of the magnitude of change, that vector looks approximately three times longer than the same vector after a Procrustes superimposition (lower half of the figure). In fact, looking at the TPS grids, or mentally joining the tips of the vectors with lines to form a triangle, clearly shows that the real change is in both cases a transformation from an equilateral triangle to an almost right triangle.

Thus, in the example using displacement vectors, the appearance of a difference in patterns, after one or the other type of superimposition, was purely an artifact of splitting a set of jointly superimposed points into subsets of landmarks. The problem may be similar adopting the within a configuration approach in studies of integration and modularity, and thus lead to potentially spurious findings. In contrast, this should not happen using the separate blocks approach on mutually exclusive modules within a structure, as each ‘module’ is re-superimposed, thus avoiding the between module covariance, which could otherwise create an appearance of integration. However, one might still wonder whether the separate blocks approach might in fact lead to the opposite issue: spuriously higher covariance within modules relative to that between them, and thus an artifactually higher degree of modularity. This is a potential issue that, although briefly discussed, will be not investigated in this paper and is left as an open question for future research.

## Simulated Examples

To explore the issue outlined in the previous section, data were created by adding isotropic normally distributed random variation (henceforth called simply ‘isotropic noise’, for brevity) around each landmark of a configuration. Since the original variation in the raw data is random, and similar in magnitude across all landmarks in that configuration, any strong pattern of variance-covariance structure in shape is purely the result of the Procrustes superimposition. Thus, also any appearance of integration or modularity will be an artifact of the Procrustes alignment used to extract shape from the original Cartesian coordinates of the raw landmarks.

2D data, and just two arbitrary ‘modules’, were used for simplicity in all analyses. However, to investigate a few different scenarios, several different configurations were employed, and sample size (N) and the amount of isotropic noise was varied across them. The number of points in each module ranged from 50:50 to ca. 3:4. Half of the datasets employed simple geometric figures (e.g., an hexagon or a circle), and the other half used configurations from previous studies on marmot mandibles (e.g.,Cardini 2003; Cardini and Tongiorgi 2003), as examples of more complex shapes and less evenly distributed landmarks.

In two datasets (the circle example and one of the mandible datasets), semilandmarks were included together with conventional landmarks. Semilandmarks are points used to discretize curves or surfaces, which lack properly corresponding landmarks (Bookstein 1997;Gunz and Mitteroecker 2013). The semilandmarks of these two configurations were either treated as conventional ‘fixed’ landmarks or slid in TPSRelw (Rohlf 2015). Sliding was achieved by minimizing either Procrustes shape distances (PRD) or bending energy (BEN) (Bookstein 1997) with five iterations. Sliding semilandmarks during the superimposition is often suggested as a way to mathematically improve the correspondence of these points across specimens in a sample. When data were analysed using the separate blocks approach, in which a configuration is subdivided and each subset of landmarks (i.e., the arbitrary module) undergoes another superimposition, slid semilandmarks were treated like normal landmarks, without sliding them again within modules. This was done to keep the analysis simple, but also because it is often said (e.g., Mitteroecker & Bookstein, 2008) that, after sliding, these points can be treated as conventional landmarks.

Configurations and modules are shown in Figure 2. The information on the number of landmarks and semilandmarks, N, the method used for sliding semilandmarks (if present), the tangent space approximation and the range of shape variation is provided in Table 1. To put the magnitude of shape variation in each dataset on a crude scale for comparisons, the mean pairwise PRD was divided by the largest possible PRD in a Procrustes shape space, which is half π. In TPSTri (Rohlf 2015), a software built to examine the properties of shape spaces, and able to simulate small, medium or large shape variation in Procrustes data using random triangles, variance is considered small, medium and large, when the mean PRD is respectively about 1%, 6% and 11% of π/2. Although arbitrary, this same convention was used here to approximately rank the variance in the simulated data of Table 1. This means that, for instance, the hexagon dataset had a large variance (ca. 10% of π/2), whereas the mandibles corresponded to small (ca. 2-3% of π/2) or medium (ca. 5% of π/2) variation depending on the dataset. For the nine landmarks mandible configurations, this type of comparison can be made more meaningful by comparing variation in simulated data with that from previous studies of real samples (Cardini, 2003; Nagorsen and Cardini 2009). This suggests that the two simulated datasets are comparable to either within species variation in North American adult marmots (mean PRD ≈ 1-2% of π/2) or interspecific differences in *Marmota* (mean PRD ≈ 4% of π/2). Thus, of the two simulated datasets, the one with smaller variance was referred to as ‘microevolutionary’ and the other one as ‘macroevolutionary’.

**Fig. 2.**
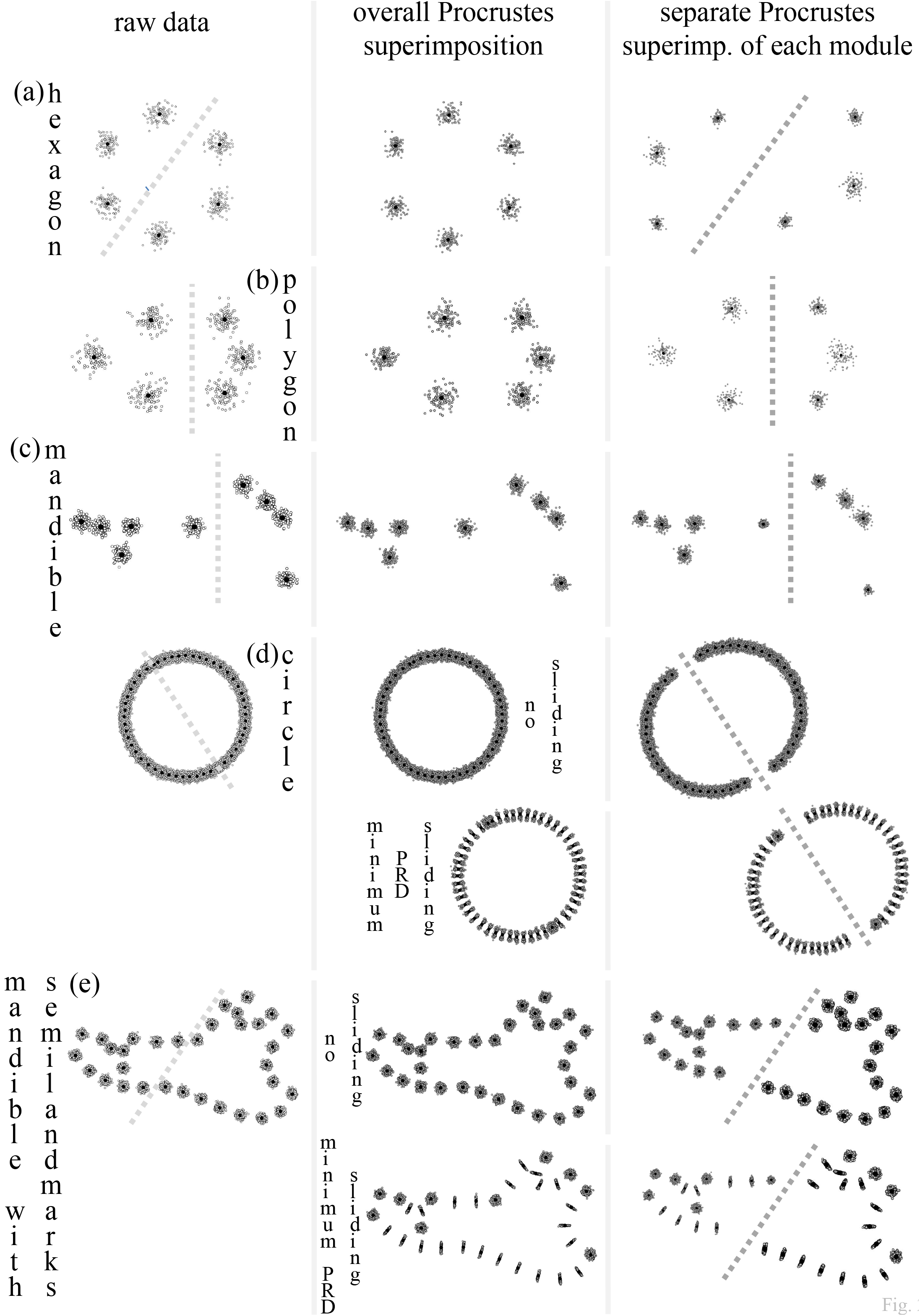
Examples of simulated datasets used in this study: left, raw data with dotted lines showing the separation into two arbitrary modules; center, common superimposition used for the within a configuration analyses of integration/modularity; right, separate superimpositions of the two modules used in the separate blocks analyses. From top to bottom: (a) hexagons; (b) six landmarks polygons; (c) marmot mandibles nine landmarks configuration (only the macroevolutionary dataset is shown); (d) circles with two landmarks (opposite extremes of the diameter) and 48 semilandmarks; (e) marmot mandibles with 10 landmarks and 23 semilandmarks. For data with semilandmarks, only configurations with semilandmarks slid with the minimum PRD criterion are shown for brevity; the minimum PRD superimposed plot also allows to clearly spot the semilandmarks, which are the sets of points approximately aligned on lines perpendicular to the outline of the study structures.

**Tab. 1.** Simulated datasets of landmarks/semilandmarks with different configurations and number of points (L), sample sizes (N) and amounts of isotropic noise. For the two configurations with semilandmarks, summary statistics are shown without any sliding and after sliding the semilandmarks using two different criteria (minimum PRD and minimum BEN). The tangent space approximation was assessed in TPSSmall (Rohlf 2015) by computing the slope and correlation in a regression through the origin of pairwise PRDs in the curved Procrustes shape space onto the corresponding Euclidean distances in the flat tangent space used for statistical analyses. The minimum, maximum and mean pairwise PRDs are shown together with the ratio of the mean PRD relative to π/2 (i.e., the largest possible PRD); using this ratio expressed as a percentage, and the conventions of TPSTri (Rohlf 2015) as an approximate guideline, variance was considered small (light grey background), medium (dark grey background) or large (black background) when respectively ≤3%, >3% but <<10%, and around 10% or more. The percentages of total shape variation explained by the first two PCs of the total configuration, as well as their ratios, are also shown; as the original raw data only contained isotropic noise, PCs should show no pattern and therefore explain approximately the same amount of variance.

| dataset | sliding method | N | L | tg space approxim |  | PRD pairwise |  |  |  | explained shape var. |  |  |
| --- | --- | --- | --- | --- | --- | --- | --- | --- | --- | --- | --- | --- |
|  |  |  |  | slope | r | min | max | mean | mean/(Pi/2) | PC1 | PC2 | PC1/PC2 |
| hexagon | none | 100 | 6 | 0.999 | 1.000 | 0.0356 | 0.2759 | 0.1511 | 10% | 17.4% | 16.0% | 1.1 |
| 6 L polygon | none | 100 | 6 | 0.998 | 1.000 | 0.0572 | 0.3827 | 0.1924 | 12% | 18.7% | 15.8% | 1.2 |
| mandibles 'microevol.' | none | 200 | 9 | 1.000 | 1.000 | 0.0160 | 0.0726 | 0.0391 | 2% | 10.7% | 9.7% | 1.1 |
| mandibles 'macroevol.' | none | 200 | 9 | 1.000 | 1.000 | 0.0284 | 0.1480 | 0.0783 | 5% | 11.0% | 9.9% | 1.1 |
| circle with<br>48 semilandmarks | none | 1000 | 50 | 1.000 | 1.000 | 0.0351 | 0.0557 | 0.0461 | 3% | 1.7% | 1.7% | 1.0 |
|  | minimum PRD | - | - | 1.000 | 1.000 | 0.0265 | 0.0680 | 0.0466 | 3% | 2.8% | 2.7% | 1.0 |
|  | minimum BEN | - | - | 1.000 | 1.000 | 0.0364 | 0.1652 | 0.0764 | 5% | 9.4% | 7.6% | 1.2 |
| mandibles with<br>23 semilandmarks | none | 1000 | 33 | 1.000 | 1.000 | 0.0293 | 0.0663 | 0.0477 | 3% | 2.4% | 2.3% | 1.0 |
|  | minimum PRD | - | - | 1.000 | 1.000 | 0.0194 | 0.0588 | 0.0381 | 2% | 3.5% | 3.4% | 1.0 |
|  | minimum BEN | - | - | 1.000 | 1.000 | 0.0239 | 0.0943 | 0.0470 | 3% | 13.6% | 7.3% | 1.9 |

## Types of Tests

The following tests of integration/modularity, based on *a priori* modules, were performed:

1) Test of integration between modules using the RV coefficient (Klingenberg 2009, 2013a), which can be seen as an extension of the Pearson correlation (r) to estimate the overall association between two blocks of multivariate variables. RV, like r, varies between zero (no integration) and one (complete integration).

2) Partial least square (PLS) test for integration using the main vector (PLS1) accounting for most covariance between modules (Rohlf and Corti 2000;Bookstein et al. 2003;Klingenberg 2013a).

3) PLS test for integration using the correlation r between PLS1 scores of the two modules. This is the same statistical framework as 2), but uses a different test statistics (which is related to, and generally in good agreement with, the test for the covariance). For this test, besides computing r, the percentage of total shape variance accounted for by PLS1 within each block was calculated. This is simply, within each module, the ratio between the variance of PLS1 scores and the sum of the variances of the shape coordinates of the landmarks in that module. These percentages are not often shown in PLS analyses, but they provide a useful information, complimentary to the percentage of covariance (test 2) and r (test 3). Indeed, one might have a vector suggesting very strong covariance and correlation, even if it accounts for a tiny proportion of total variance within modules.

4) Adams’ CR test of modularity (2016), which is a ratio of between to within modules covariances and can range between zero and more than one. This test has been proposed to provide direct evidence for modularity and overcome some of the problems with the RV coefficients (e.g., the strong effect of sample size on its estimate -Adams 2016, but see also Smilde et al. 2009and Fruciano et al. 2013). Thus, if CR is significantly smaller than 1, one can conclude that there is support for modules, as the covariance within each module is stronger than that between them.

All tests were performed using both the within a configuration and between separate blocks approaches. The only exception was the CR test, which is only available using the within a configuration method. 10000 permutations were used in each analysis to assess the significance of the test statistics. The implementations of the permutation tests are generally different depending on which approach is used (details can be found in the references provided above, as well as in the help files of the programs used for the analyses - see below). Briefly, in the between separate blocks approach, tests simply reshuffles the order of the specimens in one block at each permutation. In the within a configuration approach, each permutation works by randomly matching modules (tests 1-2-3) or randomly assigning landmarks to one or the other module without changing the number of landmarks within each of them (test 4). However, in all tests of integration (1-2-3) using the within a configuration approach, data are Procrustes re-superimposed at each round of permutation. The re-superimposition is necessary to ‘re-align’ the randomly matched modules and should take into account the additional covariance introduced by a common Procrustes superimposition (Klingenberg 2009).

Analyses were performed in MorphoJ (Klingenberg 2011) and R (R Core Team 2017) using geomorph (Adams et al. 2017) and Morpho (Schlager 2017). Some tests could only be performed in one or the other program (RV and test of significance for PLS1 covariance in MorphoJ; CR in geomorph); all other analyses, however, were done in all three programs and produced almost always totally congruent results. There were two minor exceptions of PLS1 correlations being identical in all three programs, but non-significant in MorphoJ and significant in both geomorph and Morpho; in these two cases, the congruent results of the two R programs were reported.

## Integration and/or Modularity in data generated using ‘isotropic noise’: any Significance?

Results are shown in Table 2. The tests of integration are summarized in Figure 3. RV ranged from almost zero (0.005 in the six landmarks polygon, using the separate blocks approach) to ca. 0.2, and was consistently larger in the analyses using the within a configuration approach (from 50% to more than 30 times larger). Significance was found in 12 out of 20 tests, with 10 tests being highly significant. In all instances, except two, significant RVs were found using the within a configuration approach. The two exceptions of highly significant RV using separate blocks analyses were the circles and mandibles datasets with semilandmarks slid using the minimum BEN criterion.

**Tab. 2.** Analyses of integration (1, 2, 3) and modularity (4) using different approaches (SEPARATE blocks and WITHIN a configuration) and methods: RV (1); PLS, testing both the percentage of between module covariance accounted for by PLS1 (2) and the correlation (r) between PLS1 scores in the two modules (3), with the corresponding percentages of within-module total shape variance shown in the next two columns; Adams’ (2016) CR ratio (4 - only available for the within a configuration approach). Significant tests are emphasized using a light grey background and highly signficant ones (with 0.005 threshold for high significance) using a black background. Results that contradict the expectation that the within a configuration approach might tend to suggest spurious covariance, whereas the between separate blocks method should not do it, are emphasized with bold underlined P values (e.g., mandibles with no semilandmarks showing non-significant covariance ‘despite’ a within a configuration analysis, and minimum BEN semilandmark data showing significant integration even using separate blocks).

| dataset | approach to integration/modul. | RV (1) |  | PLS1 (2) |  | PLS1 (3) |  | PLS1 % tot. variance |  | modularity test (4) |  |
| --- | --- | --- | --- | --- | --- | --- | --- | --- | --- | --- | --- |
|  |  | value | P | % covar. | P | r | P | module 1 | module 2 | CR | P |
| hexagon | SEPARATE | 0.015 | 0.53 | 100% | 0.47 | 0.175 | 0.47 | 55% | 47% | - | - |
|  | WITHIN | 0.209 | 0.0001 | 38% | 0 | 0.735 | 0.0001 | 17% | 20% | 0.940 | 0.1946 |
| 6 L polygon | SEPARATE | 0.005 | 0.9102 | 99% | 0.8905 | 0.100 | 0.8881 | 54% | 46% | - | - |
|  | WITHIN | 0.152 | 0.0003 | 40% | 0.0046 | 0.756 | 0.0095 | 27% | 9% | 0.767 | 0.0958 |
| mandibles 'microevol.' | SEPARATE | 0.030 | 0.2 | 50% | 0.33 | 0.272 | 0.29 | 18% | 23% | - | - |
|  | WITHIN | 0.067 | 0.01 | 32% | 0.09 | 0.423 | 0 | 14% | 13% | 0.558 | 0.0086 |
| mandibles 'macroevol.' | SEPARATE | 0.028 | 0.25 | 48% | 0.45 | 0.232 | 0.59 | 17% | 31% | - | - |
|  | WITHIN | 0.063 | 0.03 | 31% | 0.21 | 0.520 | 0 | 14% | 8% | 0.531 | 0.0086 |
| circle with semilandmarks | SEPARATE | 0.041 | 0.98 | 9% | 0.28 | 0.373 | 0.67 | 3% | 2% | - | - |
|  | WITHIN | 0.056 | 0.0001 | 9% | 0.0001 | 0.620 | 0.0001 | 1% | 2% | 0.894 | 0.0001 |
| slid minimum PRD | SEPARATE | 0.023 | 0.92 | 14% | 0.87 | 0.276 | 0.73 | 4% | 4% | - | - |
|  | WITHIN | 0.038 | 0.0001 | 22% | 0.0001 | 0.721 | 0.0001 | 3% | 2% | 0.384 | 0.0001 |
| slid minimum BEN | SEPARATE | 0.019 | 0.0010 | 31% | 0.0400 | 0.256 | 0.0001 | 7% | 10% | - | - |
|  | WITHIN | 0.146 | 0.0001 | 48% | 0.0001 | 0.708 | 0 | 10% | 9% | 0.456 | 0.0001 |
| mandibles with semilandmarks | SEPARATE | 0.025 | 0.97 | 15% | 0.41 | 0.308 | 0.41 | 5% | 3% | - | - |
|  | WITHIN | 0.040 | 0.0001 | 14% | 0.0001 | 0.651 | 0.0001 | 2% | 3% | 0.721 | 0.0001 |
| slid minimum PRD | SEPARATE | 0.018 | 0.74 | 20% | 0.41 | 0.251 | 0.33 | 6% | 5% | - | - |
|  | WITHIN | 0.034 | 0.0001 | 20% | 0.0001 | 0.772 | 0.0001 | 2% | 4% | 0.414 | 0.0001 |
| slid minimum BEN | SEPARATE | 0.113 | 0.0001 | 82% | 0.0001 | 0.599 | 0.0001 | 17% | 11% | - | - |
|  | WITHIN | 0.225 | 0.0001 | 74% | 0.0001 | 0.744 | 0 | 13% | 16% | 0.640 | 0.0001 |

**Fig. 3.**
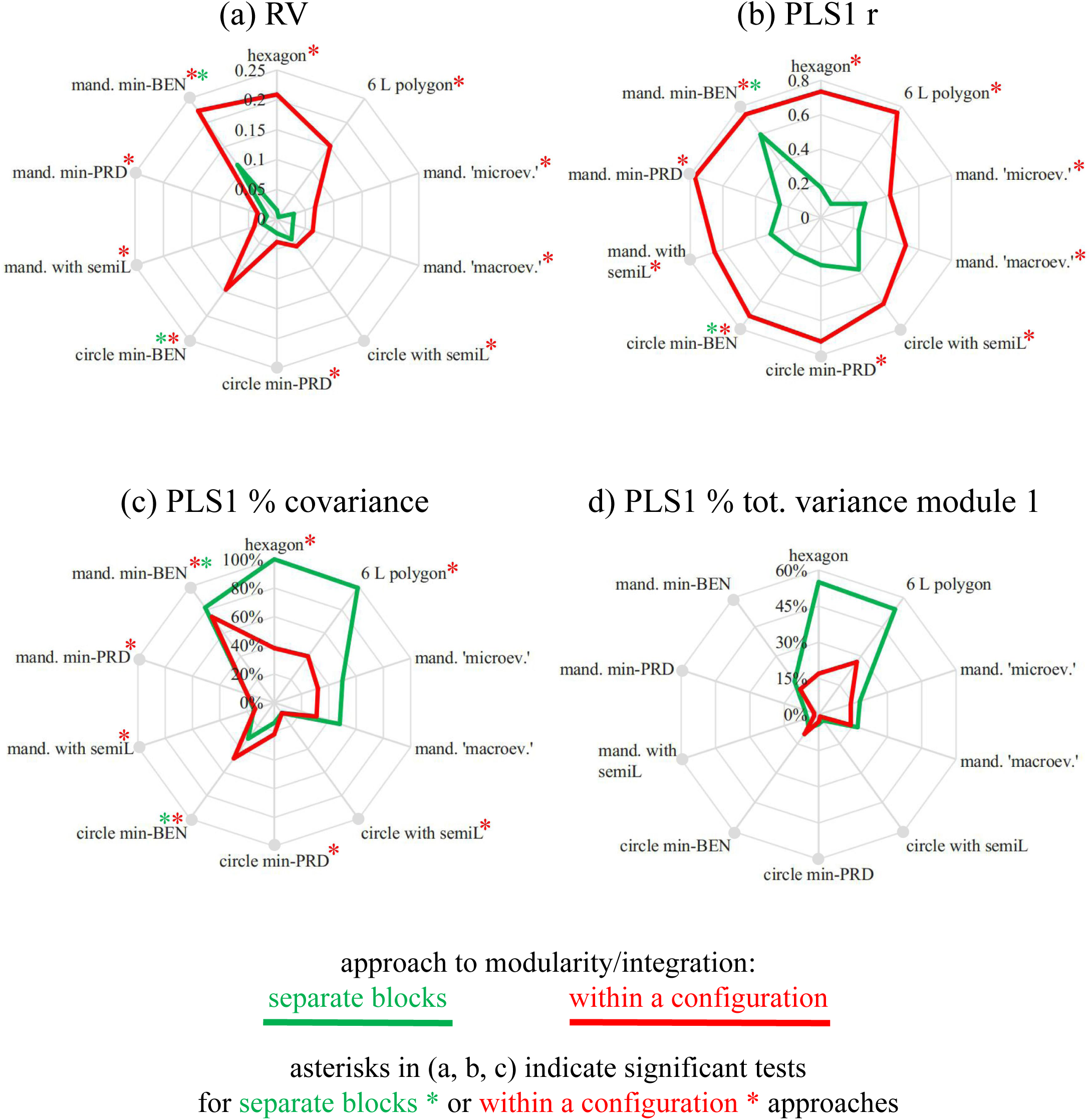
Spider plots showing for all datasets: (a) RV coefficients, (b) correlations r between PLS1 scores of the two modules, (c) percentages of shape covariance explained by PLS1 and (d) percentages of total shape variance explained by PLS1 for the first module (that of the second module is not shown, as the pattern of differences between the two approaches is almost identical). Green lines are used to connect values obtained with the separate blocks approach and red ones for the within a configuration approach, and asterisks indicate signficant tests in (a, b, c) using one or the other approach depending on the colour; lines ending with grey circles emphasize datasets with semilandmarks.

The PLS analysis was in good agreement with the RV tests. I describe first the results for the correlation of PLS1 scores, because they were almost perfectly congruent with those using RV: r ranged from ca. 0.1 to 0.8, with r from 20% larger to up to 8 times bigger in analyses using the within a configuration approach; overall, 12 of the 20 tests were significant (11 of them highly significant), and these were, as with RV, all those using the within a configuration approach plus the separate blocks analyses of the two minimum BEN slid configurations.

The tests for the covariance accounted for by PLS1 were also mostly in good agreement with both RV and r tests, as significant analyses involved precisely the same datasets as in those analyses, except the two mandible samples (‘micro-’ and ‘macroevolutionary’, with no semilandmarks). For these two datasets, none of the tests, regardless of the approach, reached significance. Both the percentages of covariance, as well as those of variance accounted for by PLS1, were generally smaller using the within a configuration approach, with the main few exceptions being some of the slid semilandmarks analyses.

Overall, the vast majority of the tests for integration suggested a strong covariation between modules but only when using the within a configuration approach. Using the between separate blocks approach, despite PLS1 vectors accounting for generally more covariance and variance, none of the tests supported integrated modules, except the two minimum BEN slid semilandmark datasets.

Interestingly, when datasets were tested for modularity using CR and the within a configuration approach, all CRs were smaller than one (range: 0.4-0.9), and most of them (with the exception of the two smallest configurations, the hexagon and six landmarks polygon) were significantly smaller than expected by randomly assigning landmarks to partitions. Thus, because CR<1 indicates more covariance within modules compared to between modules, these results supported modularity in eight of the 10 analyses. As the CR test is not available using the separate blocks approach, the two approaches could not be compared and this is why the total number of tests was just 10.

## Point-Estimates *VS* Simulations

All results reported in the previous section represent point-estimates from single analyses using several different test statistics and two general approaches (within a configuration or separate blocks) on 10 different sets of data (considering, for the sake of brevity, the slid datasets as different types of data). The vast majority of these 10 sets of data, when analysed using the within a configuration approach, produced congruent results indicating a serious issue with type I errors (i.e., obtaining a false postive by rejecting the null hypothesis when this is in fact the truth). It seems unlikely that so many instances of false positives from different analyses and datasets could be found by chance but one might wonder what would happen if the experiments using random noise were repeated many times on the same data. Although this goes beyond the stated aim of this study (i.e., raising an issue based on preliminary evidence), thanks to an R script made available by an anonymous reviewer, type I error rates were briefly explored in all datasets without sliding semilandmarks. This was done by running 100 times PLS tests for the first pair of vectors using both approaches, as well as CR tests using the within a configuration approach. In all datasets, the amount of random isotropic noise added to the data in each run of the simulations was approximately the same as in the point-estimate analyses (as assessed by the mean and maximum pairwise Procrustes distance in a sample). The number of specimens in the simulated samples was equal to 100 except for the circles and mandibles with unslid semilandmarks, where it was increased to 500 to take into account the much larger number of variables of these two sets of data. Table 3 reports the resulting estimates of type I error rates, which are the proportion of times significance was found. These should not exceed 0.05, the nominal threshold usually adopted for type I errors. Congruently with the point-estimates of the previous section, the rate of false positives ranged from 0.01 to 0.09 for PLS1 analysed with separate separate blocks. In contrast, using the within a configuration approach, both PLS1 and CR showed a much higher rate of type I errors, ranging from 0.27 (ca. 5 times the nominal value) to 1.00 (i.e., 100% false positives). Thus, also this small set of simple simulations strongly supported the outcome of the point-estimates study: the separate blocks approach seems appropriate in terms of type I error rates, whereas the within a configuration analyses largely inflates the occurrence of false positives.

**Tab. 3.** Type I error rate (i.e., proportion of true null hypotheses incorrectly rejected) for PLS1 and CR tests estimated in 100 simulations using N=100 for all datasets except the circle and the mandible with unslid semilandmarks, whose N=500. Rates equal to or larger than 0.1 are emphasized with a black background.

| dataset | approach to integration/modul. | PLS1 | modularity test CR |
| --- | --- | --- | --- |
| hexagon | SEPARATE<br>WITHIN | 0.03<br>1.00 | -<br>0.27 |
| 6 L polygon | SEPARATE<br>WITHIN | 0.01<br>1.00 | -<br>0.50 |
| mandibles 'microevol.' | SEPARATE<br>WITHIN | 0.09<br>0.54 | -<br>1.00 |
| mandibles 'macroevol.' | SEPARATE<br>WITHIN | 0.07<br>0.55 | -<br>1.00 |
| circle with semilandmarks | SEPARATE<br>WITHIN | 0.06<br>0.90 | -<br>0.99 |
| mandibles with semilandmarks<br>(no sliding) | SEPARATE<br>WITHIN | 0.07<br>0.99 | -<br>1.00 |

## Interpretations of Main Patterns

Before discussing the main results, it is important to stress that the datasets and types of tests used in this paper were not aimed at thoroughly assessing the statistical properties of the methods. They are examples, analysed with common methods and used to explore whether there is a problem. If indeed they suggest a potential issue, that will require extensive studies to assess its importance and generality using simulations and a large number of different scenarios (e.g., landmark number and density, modules with large differences in number of landmarks, three or more modules, different proportions of landmarks and semilandmarks, a variety of sample sizes, different amounts of ‘real’ covariance etc.).

It is also useful to emphasize that P values were not corrected for multiple testing and, more importantly, that they are used in this context mainly as a numerical aid to better appreciate possible misleading results from analyses conducted in some of most widely used programs for Procrustean geometric morphometrics. In general, P values should be used and interpreted with caution, and it has been suggested that in morphometrics (Bookstein 2017), as in other fields, significance testing should not be an aim in itself. Also, as an important cautionary note in interpreting significant tests in this study, readers should bear in mind that simulated samples were large in relation to the number of variables, thus increasing power. Having large samples is desirable but it might have overemphasized the importance of small covariances introduced by the superimposition: in true biological data, where real covariance is expected and might be much larger than that due to the superimposition, the problem of spurious results might be less concerning. However, even in that case, the superimposition (including, where appropriate, the sliding of semilandmarks) will alter the variance-covariance structure of the data, and the extent to which this might affect results will be probably hard to assess. Also, in terms of power and sample size, as recently stressed by Bookstein (2017), one should bear in mind that an unfavourable ratio between the number of specimens and the number of variables can lead to inaccurate findings. For instance, if the type I error rate simulations using the two configurations with more points (mandibles and circles with, respectively, 33 and 50 points, and thus ca. 60 to almost 100 shape variables) were run using N=100 (instead of 500, as in Tab. 3), the proportion of false positives using the within a configuration approach would decrease to 0.07-0.11 in all tests except the mandible CR (that would be large - 0.53 - but, nevertheless, almost half than found using N=500). Thus, even when using the within a configuration approach, with an inadequate sample size compared to the large number of variables, some simulations might have misleadingly suggested no problem (or a minor one). In fact, this applies also to the three datasets (pupfish, hummingbirds and scallops, all of them available as examples in geomorph -Adams et al. 2017) used by the reviewer in her/his original simulations: using an unwisely small sample size (N=100), in relation to data dimensionality (from almost 50 to more than 130 shape variables in those three configurations), type I errors seem appropriate; however, by changing a single parameter in the R script, that is using N=500, type I error rates become hugely inflated, ranging from 0.33 (hummingbirds) to 0.71-0.96 (scallops and pupfish, respectively).

The general pattern suggested by the analyses of example datasets generated using isotropic noise, as well as the corresponding small set of simulations for estimating type I error rates, is clear: using raw data that do not contain any covariance, within or between *a priori* arbitrary modules, some of the most common methods used for the analysis of integration and modularity of Procrustes shape coordinates can lead to spurious conclusions. As expected, this is particularly evident for the within a configuration approach. With this approach, the covariance found between arbitrary modules from an isotropic noise model of variation is simply a by-product of the Procrustes superimposition. This artifactual between modules covariance, that can lead to the false appearance of integration, is generally ‘removed’ using the separate blocks analysis, in which each module undergoes a separate superimposition. However, separate Procrustes superimpositions will still increase covariance within each module, whereas they might potentially remove any real covariation between modules related to size and reciprocal orientation, therefore underestimating integration and potentially overestimating modularity. Thus, in tests of integration, the separate blocks approach may have appropriate rates of type I errors (no more false positives than expected by chance because of sampling error), but yet lead to large type II error rates (i.e., too many false negatives suggesting no integration when in fact it is there). As anticipated, this is an aspect not examined in this paper, but certainly important in future studies on the effects of the superimposition in tests of modularity and integration.

The superimposition may also explain why, using the within a configuration approach, one can find contradictory evidence of both integration and modularity. This is because the covariance introduced by a common superimposition is ‘spread’ across the whole configuration but is also likely to be stronger between contiguous landmarks (i.e., within a ‘module’). This is more easily appreciated using an example. One could replicate the analyses using the within a configuration approach on non-sensical ‘modules’ made by selecting every other landmark/semilandmark in one module, and the remaining points in the other. If this was done, for instance using the nine landmarks mandibular macroevolutionary dataset or the unslid circles, the tests for integration (1-2-3) would remain highly significant in both cases (P<0.005), but CR would become larger than one (1.1-1.2) and no longer significant. This is because, in these ‘every-other-point modules’, the ‘contiguity effect’ of the superimposition is destroyed (thus reducing the apparent within module covariance), whereas the large scale covariation across all landmarks is still there to create spurious findings of integration.

Another apparently puzzling result is that, in the PLS analyses, the separate blocks approach tends to explain more covariance, and within module variance, but the corresponding PLS1 vectors are almost always non-significant. In fact, however, the observation of higher covariance and variance of PLS1 using the separate blocks approach is most pronounced in the configurations with fewer points (i.e., all those without semilandmarks). This is because the separate superimpositions of the two modules lead to a loss of more degrees of freedom than a single common superimposition, as in each GPA size, translational (along the X and Y axes, in 2D) and rotational variation are removed. Thus, the real number of informative dimensions is smaller with separate blocks, and that might somewhat inflate the amount of covariance and variance accounted for by the main direction of covariation between modules, something that becomes especially evident with less landmarks and, therefore, less shape variables to start with.

Much more interesting is that even the separate blocks approach can produce spurious evidence for integration in special cases: these, in the examples used in this sudy, were the datasets where semilandmarks were slid using the minimum bending energy criterion. This step in the superimposition introduces a degree of covariation across the points such that its effect, at least when the number of semilandmarks is large compared to that of ‘fixed’ landmarks, is felt even if modules are later resuperimposed on their own. To control for this, one might try to re-slide the semilandmarks in the separate blocks, a step that was not included in this study.

In general, morphometricians should be particularly careful not to treat slid semilandmaks as ‘conventional landmarks’, as not only they lack the precise anatomical correspondence of ‘fixed’ landmarks (Klingenberg 2008; Oxnard and O’Higgins, 2009; Cardini 2013; Cardini and Loy 2013), but they are also affected by the mathematical treatment used to increase their geometric correspondence. That sliding can introduce patterns that were not in the original data can be seen also using a simple PCA. Figure 4 shows, using the mandibles with semilandmarks as an example, scatterplots of the first two PCs and bar plots for the percentages of variance accounted for by each vector. With isotropic noise, scatterplots should show circular variation around the mean and PCs explain approximately the same amount of variance. Indeed, this is reasonably true for the Procrustes shape coordinates without sliding and using the minimum PRD sliding method (Fig. 4a-b), which reduces some of the random variation in the semilandmarks by sliding them along an estimated tangent (approximated at each point by a chord between its adjacent points), so that it becomes mostly orthogonal to the tangent and does not tend to dominate any PC axis. In contrast, using the minimum BEN criterion, which retains some of that tangential variation and thus creates a pattern of ‘quasi-oval’ scatter around the mean position of each semilandmark, PC1 becomes elongated and accounts for twice more variance than PC2 and 4-5 times more than PC1 without sliding or sliding using the minimum PRD method.

**Fig. 4.**
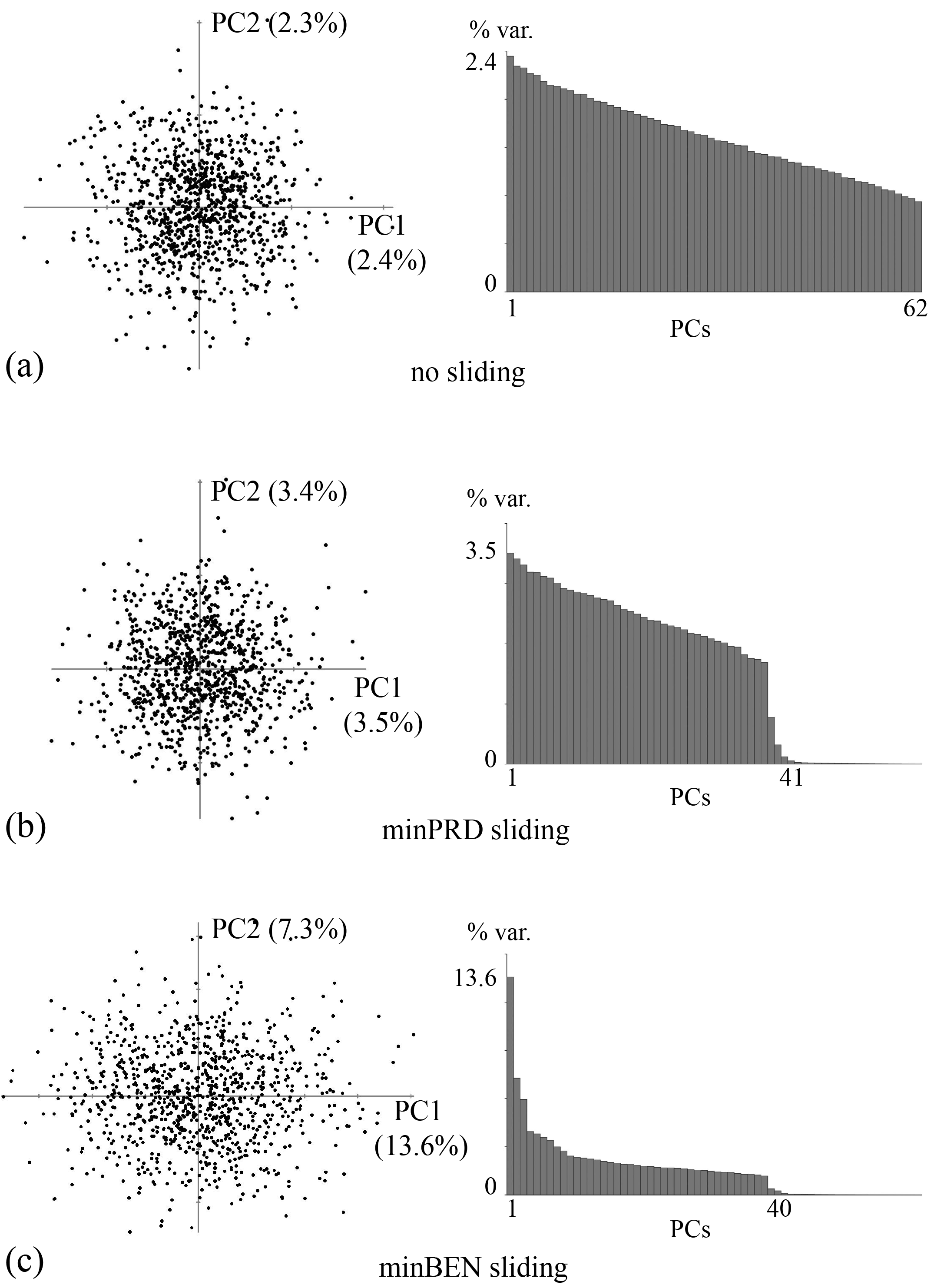
PCAs of mandibles with semilandmarks: the scatterplots of the first two PCs are shown on the left, and bar plots for proportions of variance accounted for by each PCs are shown on the right; (a) Procrustes superimposed data with no sliding; (b) semilandmarks slid using minimum PRD; (c) semilandmarks slid using minimum BEN.

Although in relation to a different issue, that of an unfavourable ratio between the number of variables and sample size, a similar problem of disparity in variance accounted for by PCs of random normally distributed data has been recently brought to the attention of the morphometric community by Bookstein (2017). It seems that sliding using the minimum BEN criterion could aggravate this problem, and worryingly that is likely to happen in cases where one already has many variables and relatively small samples, as it is not unusual in analyses of outlines or surfaces using semilandmarks. For instance, in a very recent investigation of mosaic evolution in the avian cranium, Felice and Goswami (2018) analysed ca. 350 species of birds employing about 30 landmarks and almost 300 semilandmarks, a large number of points that should lead to “not simply a statistical artifact, but rather … [to a] better representation of morphology and the relationships among different aspects of shape” (p. 2, SI Appendix). That more data are obtained this way is certainly true, but one can also wonder to what extent patterns of covariance, as well as the comparison of models with different modules, and the estimates of within-module total variance, might have been influenced by the common superimposition, including the minimum bending energy sliding of ca. 10 times more points than the number of ‘fixed’ landmarks. This does not mean that findings in that study must be flawed, but it urges to be cautious before concluding that more is inevitably better than less.

## Conclusions

In this preliminary investigation, I have explored whether the commonly used superimposition procedures necessary to extract shape variables from Cartesian coordinates of landmarks might introduce artifactual patterns of variance-covariance that can lead to spurious results. This has been investigated in the context of analyses of integration and modularity, but it may concern other types of studies in which subsets of landmarks from a common superimposition are analysed and compared. The problem is not dissimilar to that of (over-)interpreting displacement vectors in the visualization of shape differences (as exemplified in Viscosi and Cardini 2011).

Bearing in mind its exploratory nature, the limited number of tests examined and a main focus on type I errors, and therefore the need of further in depth research considering a large variety of scenarios as well as type II errors, three main messages can be taken from this study, that might hopefully stimulate future investigations, as well as recommend a degree of caution in applications of Procrustes methods:

1) The superimposition can introduce an amount of covariance (Rohlf and Slice 1990;Mitteroecker and Bookstein 2007;Klingenberg 2009;Adams 2016) that can lead to spurious results in testing integration and modularity. This can be especially problematic using the within a configuration approach and may happen even if the test recomputes the superimposition at each randomization (Klingenberg 2009), as shown by finding significance from data containing only random isotropic noise. This does not mean that users must necessarily prefer the separate blocks over the within a configuration approach: they may have different aims and issues (Baab 2013;Klingenberg 2009, 2013), but, especially if one finds that they lead to different conclusions on the same data, a morphometrician should be extremely cautious in interpreting results. Indeed, even if separate blocks may be less affected by the risk of spurious results at least in the tests of integration (and this may not be true for the tests of modularity!), as pointed out by Adams in MORPHMET, this approach excludes potentially important information (i.e., relative size and position of the modules) that, as suggested by a reviewer, might lead to “higher type II error than a single GPA, as portions of covariation between modules will not be properly preserved under separate GPA approaches”.

2) The ‘contiguity effect’ induced by a common superimposition might explain ambiguous results suggesting both integration and modularity. This is definitely an issue that requires more study to be better understood, and might partly depend also on different permutation schemes employed to test different hypotheses. It is also not impossible that this effect, as well as the more general ones on variance-covariance patterns discussed in the previous point, may have been overemphasized in large simulated datasets purely made of isotropic variation. Yet, one cannot be sure, and indeed the impact of these issues may vary from case to case and generalizations might be difficult to make.

3) Even after, and probably especially after, sliding them, semilandmarks cannot be “used in … geometric morphometrics as if they were homologous point locations” (Mitteroecker and Bookstein 2008, p. 946). From a theoretical point of view, answering the question of the biological correspondence of semilandmarks is far from trivial (Klingenberg 2008;MacLeod 2008; Oxnard and O’Higgins 2009). In practice, it seems that sliding might contribute to increase covariance and in this respect the minimum BEN criterion could sometimes be more problematic. Clearly, this does not mean that one should never use semilandmarks or inevitably prefer minimum PRD to minimum BEN for sliding. However, it does suggest that there is no garantee that more points necessarily lead to increased accuracy and, if and when semilandmarks are really crucial, one should also acknowledge their limitations and potential issues (including those related to the large number of variables they generate in relation to sample size -Bookstein 2017).

As a concluding paragraph, it is worth mentioning that Bookstein (2015)has recently proposed a new interesting approach to explore integration and modularity, that breaks this methodological dichotomy by bringing both types of analyses together in a single continuum. As in other exploratory methods (Goswami and Polly 2010), his method neither employs a test or requires *a priori* modules. Instead, it looks for relationships between the variance of partial warps and bending energy. Put it informally, partial warps provide a way of extracting from landmark coordinates shape variables that capture variation at different spatial scale (Bookstein 1991;Rohlf 1998), from changes occurring almost uniformly over the whole landmark configuration (thus, requiring the least BEN) to those highly localized (therefore, needing large BEN). The method has some promising aspects since, among other things, by using only partial warps, it should not be affected by the choice of superimposition. This is because the purely uniform variation (e.g., dilation, stretching or shearing occurring in exactly the same way over the entire landmark configuration) is omitted, although by, doing so, it also excludes a potentially interesting aspect of integration. Nevertheless, Bookstein (2015)suggested to plot, after logarithmic transformations, the variance of partial warps onto their corresponding BEN. Then, the slope of a regression line approximating the data will be approximately zero for isotropic noise, −1 for self-similarity across all spatial scales, between zero and −1 for ‘disintegrated’ change (i.e., no integration and thus potential modularity), and more negative than −1 for global integration. Thus, as an example, I applied this approach to the mandible dataset with semilandmarks (results - not shown - were similar using circles, but less interesting in the other datasets, that have too few landmarks for producing interpretable patterns). The method performed (Fig. 5) as expected for unslid data and data with semilandmarks slid using the minimum PRD criterion. In both cases, the slope was almost zero, as it should for isotropic data. This means that contrary to other within a configuration methods, such as RV, PLS and CR, Bookstein’s ‘integration-disintegration’ technique did not misleadingly suggest integration or modularity in random landmarks with isotropic variance. However, when applied after sliding with the minimum BEN criterion, the slope was approximately −0.4, which indicates ‘disintegration’ but, as for the finding of a significant CR, and probably for the same reason (‘contiguity effect’), the result is misleading and thus shows again that sliding does not ‘fix’ semilandmarks and that these points are indeed ‘special’.

**Fig. 5.**
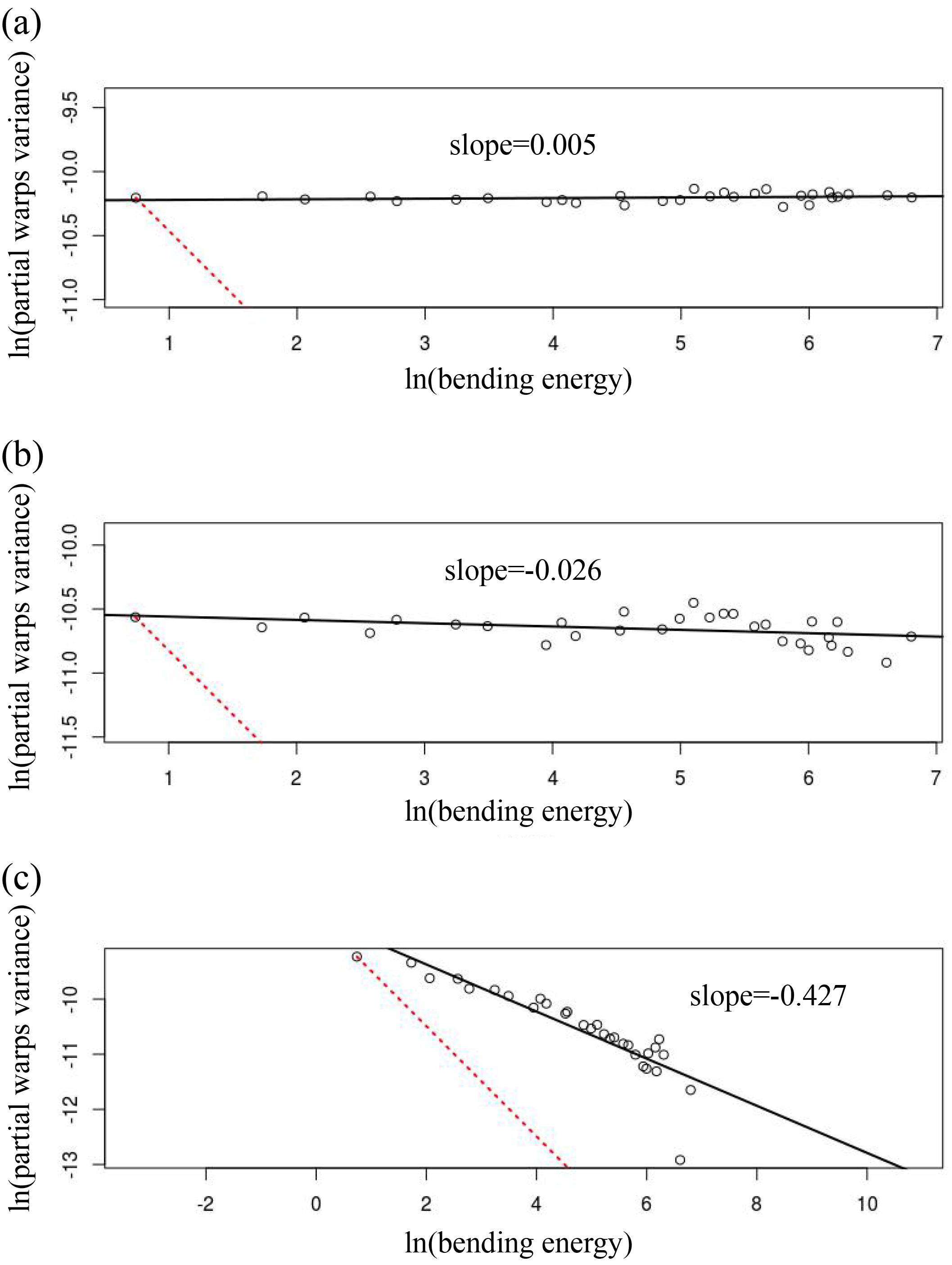
Bookstein (2015) ‘integration-disintegration’ approach on simulated mandibles with semilandmarks: regression line (solid) of partial warps variance *vs* bending energy (log-transformed), with the dotted line showing the expectation for self similarity (slope=−1); (a) Procrustes superimposed data with no sliding; (b) semilandmarks slid using minimum PRD; (c) semilandmarks slid using minimum BEN.

## ACKNOWLEDGEMENTS

I am deeply grateful to Paul O‘Higgins for critically discussing with me some of the points presented in this communication, and to Dean Adams for his fundamental help and explanations on some of the methods commonly used in the analysis of modularity and integration. I owe a huge thank to Jim Rohlf, for reading the quasi-final version of this paper and for being most supportive and constructive in his assessment of this work. I am also greatly in debt to David Polly and two anonymous reviewers for their in depth assessment of a previous version of this paper, including DP and another reviewer exploring the issues further using simulations: regardless of agreement or disagreement, all their comments and suggestions greatly improved this preliminary investigation and contributed to showing that there is an issue and that this is definitely more complicated that I originally thought!

Last but not least, I wish to dedicate this paper to the memory of Paolo Tongiorgi (1936-2018): Paolo, you have been an extraordinary scientist; an incredibly supportive mentor; a wonderful friend; and the greatest example of generosity I have known in my academic career.

